# CHLOROPLAST GENOME AND PHYLOGENETIC ANALYSIS OF KATMON (*Dillenia philippinensis* Rolfe), A PHILIPPINE ENDEMIC FRUIT

**DOI:** 10.1101/2025.11.26.690882

**Authors:** Jairus Jake M. Lucero, Judy Ann M. Muñoz, Lyka Y. Aglibot, Don Emanuel M. Cardona, Lavernee S. Gueco, Aprill P. Manalang, Jeric C. Villanueva, Roneil Christian S. Alonday

## Abstract

**Background:** Katmon (*Dillenia philippinensis* Rolfe) is a Philippine endemic fruit species with a relatively well-studied biochemical profile but poor genomic characterization. Studies involving the chloroplast genome can provide valuable insights into its evolution and support conservation efforts.

**Methods:** The complete chloroplast genome of *D. philippinensis* was sequenced using Illumina NovaSeqX. Reads were quality-checked, assembled with GetOrganelle, and annotated using CPGAVAS2 and GeSeq. Simple sequence repeats, codon usage, and inverted repeat boundaries were analyzed. Phylogenetic relationships were inferred using concatenated *rbcL* and *matK* sequences via maximum likelihood analysis.

**Result:** The chloroplast genome was 161,591 bp with a GC content of 36.3%. It exhibited the typical quadripartite structure, consisting of a large single-copy region (89,411 bp), a small single-copy region (19,208 bp), and a pair of inverted repeats (26,486 bp each). A total of 113 unique genes were identified, comprising 79 protein-coding, 30 tRNA, and four rRNA genes. Fifty-four SSRs, primarily A/T mononucleotide repeats, and 53,863 codons were observed. Phylogenetic analysis placed *D. philippinensis* as the closest relative to *D. suffroticosa* and the most distantly related to *D. ovata*. The complete chloroplast genome of *D. philippinensis* provides a valuable resource for phylogenetic studies, germplasm characterization, and future breeding and conservation programs.

## INTRODUCTION

Endemic plants are species that are solely found within a specific geographical region. They are well-adapted to their local habitats, thereby posing less damage to the environment and making them valuable resources for reinforcing food security. They also exhibit greater climate resiliency and pest tolerance than commercial crops (Nhamo *et al*., 2022). Moreover, indigenous fruits display substantial, if not higher, nutritional value compared to widely cultivated species (Durst and Bayasgalanbat, 2014). Despite the health benefits and potential for diet diversification, indigenous plants remain poorly studied (Oraye *et al*., 2023; Villarino and Villarino, 2023), leading to their underutilization and preventing their full potential from being harnessed.

*Dillenia philippinensis* is a medium-sized, evergreen fruit tree belonging to the family *Dilleniaceae* (Aquino *et al*., 2015, as cited by Fatallo and Panes, 2022). It is distributed in many provinces across the Philippines, such as Laguna, Quezon, Oriental Mindoro, and Cebu (Magdalita *et al.,* 2014). The leaves of this tree species were found to exhibit cytotoxic (Dante *et al*., 2019), antifungal (Ragasa *et al*., 2009), and antioxidant (Ansari *et al*., 2021) activities. Meanwhile, the fruit extracts display antimicrobial activity (Tubillo *et al*., 2016) and can be used as a natural food preservative (Pormento, 2024). While the biochemical profile of katmon is relatively well-studied, its genomic characteristics remain largely unexplored. Genomic characterization is limited to the work of Fatallo and Panes (2022), who used rbcL, matK, and ITS markers for barcoding and phylogenetic analysis. No further studies have been conducted for the genetic characterization of this species.

The chloroplast genome is one of the three genomes present in plants (Rozov *et al.,* 2022). It holds evolutionary significance, as it is absent in other eukaryotes and functions for photosynthesis, a process whose biochemical mechanisms are conserved among plants (Theeuwen *et al.,* 2022). Thus, the characterization of a chloroplast genome provides valuable insights into phylogenetic relationships and plant taxonomy (Daniell *et al.,* 2016). Moreover, a comprehensive analysis of the chloroplast genome can equip plant breeders and conservation biologists with pertinent knowledge on breeding, conservation, and germplasm characterization efforts. Molecular data derived from the chloroplast genome can support the development of new cultivars with greater yield, higher nutritional content, and higher genetic diversity (Swarup *et al.,* 2020). Therefore, this study sought to characterize the chloroplast genome of *D. philippinensis* to gain a deeper understanding of its phylogeny.

## MATERIALS AND METHODS

### Sample Collection

Mature leaves of *D. philippinensis* accessions GB68492 and GB70976 were collected from the germplasm collections of the National Plant Genetic Resources Laboratory, Institute of Plant Breeding, University of the Philippines, Los Baños. The collected leaves were placed inside airtight plastic bags and kept in an icebox during transportation. The leaves were then disinfected using 70% ethanol, put in a resealable plastic bag, and stored at -80°C until further use.

### Genomic DNA Isolation

Genomic DNA was extracted from collected leaf samples following the protocol of Inglis *et al*. (2018). The purity and concentration of the isolated DNA were determined using an Epoch Multi-Volume Spectrophotometer System (BioTek Instruments Inc., USA). The extract from accession GB68492 was sent to Macrogen (South Korea) for complete chloroplast genome sequencing via the Illumina NovaSeqX platform (Illumina Inc., San Diego, CA).

### Chloroplast Genome Assembly and Annotation

Raw reads generated from Illumina sequencing were subjected to quality checking using FastQC v0.12.1 (Andrews 2010). Adapters and low-quality reads were trimmed using Trimmomatic v0.39 (Bolger et al., 2014), which was implemented using the following operations: “ILLUMINACLIP TruSeq3 2:30:10:8; LEADING:25; and SLIDINGWINDOW:4:20”. Poor-quality reads were discarded, while high-quality reads were subjected to de novo assembly using GetOrganelle v1.7.7.1+ (Jin et al., 2020), with the consensus rbcL sequence serving as a seed sequence for assembly. The assembled chloroplast genome was annotated using CPGAVAS2 (Shi et al., 2019), with the complete chloroplast genome of D. indica deposited in NCBI GenBank (NC_042740.1) serving as reference. Manual validation was done using Geneious Prime v2025.1.2 and by cross-checking the annotations made by GeSeq (Tillich et al., 2017).

### Amplification of rbcL and matK

The genomic DNA samples were diluted to 20 ng/μL. Polymerase chain reaction (PCR) was performed to amplify rbcL and matK. Primers from Thooptianrat *et al*. (2017) and Yu *et al*. (2011) were used to amplify rbcL and matK, respectively. PCR thermocycling conditions were adapted from the study of Fatallo & Panes, 2022. The amplicons were electrophoresed using 1.5% agarose at 110 V for 30 min to check amplification success. They were then purified using a NucleoSpinTM Gel and PCR Clean-Up Kit (Macherey-Nagel, Germany). Spectrophotometry was performed to assess the purity and concentration of the purified amplicons. Lastly, the amplicons for rbcL and matK were sent to Macrogen (South Korea) for bidirectional Sanger sequencing.

### Phylogenetic Analysis

The consensus *rbcL* and *matK* sequences of the two *D. philippinensis accessions* were subjected to phylogenetic analysis to confirm their relationship with other species in the family Dilleniaceae. The reference rbcL and matK sequences of each species were obtained from NCBI GenBank. Jojoba (Simmondsia chinensis) (NC_040935) and pokeweed (Phytolacca americana) (NC_067846) were used as outgroups. Multiple sequence alignment of the *rbcL* and *matK* sequences was performed using ClustalW (Thompson *et al*., 1994). The aligned *rbcL* and *matK* sequences were also concatenated using FaBox v.1.61 (Villesen 2007). Finally, phylogenetic trees were constructed based on the aligned *rbcL*+*<u>matK</u>*sequences, with best-fit substitution models of Tamura 3-parameter model with gamma distribution (T92 + G), Kimura 2-parameter model with gamma distribution (K2 + G), and Tamura 3 parameter model with gamma distribution (T92 + G), respectively. The phylogenetic trees were constructed using MEGA12 (Kumar *et al*., 2024), following the maximum likelihood method with the respective best-fit substitution model and employing bootstrap replicates of 1000.

## RESULTS AND DISCUSSION

### Chloroplast Genome Assembly and Annotation

The complete chloroplast genome of *Dillenia philippinensis is* 161,591 bp in length with 36.3% GC content and exhibits the typical quadripartite structure composed of a large single copy (LSC, 89,411 bp), a small single copy (SSC, 19,208 bp), and two inverted repeat (IRa and IRb) regions (26,486 bp each) (Figure 1). A total of 130 genes were identified (Table 1), comprising 113 unique genes and 16 duplicated in the IRs. Duplicated genes include five protein-coding (rpl2, rpl23, ycf2, ndhB, rps7), seven tRNA (trnI-CAU, trnL-CAA, trnV-GAU, trnA-UGC, trnR-ACG, trnN-GUU), and four rRNA genes (rrn23S, rrn4.5S, rrn5S, rrn16S). The trans-spliced rps12 gene was also annotated twice.

**Figure 1.**
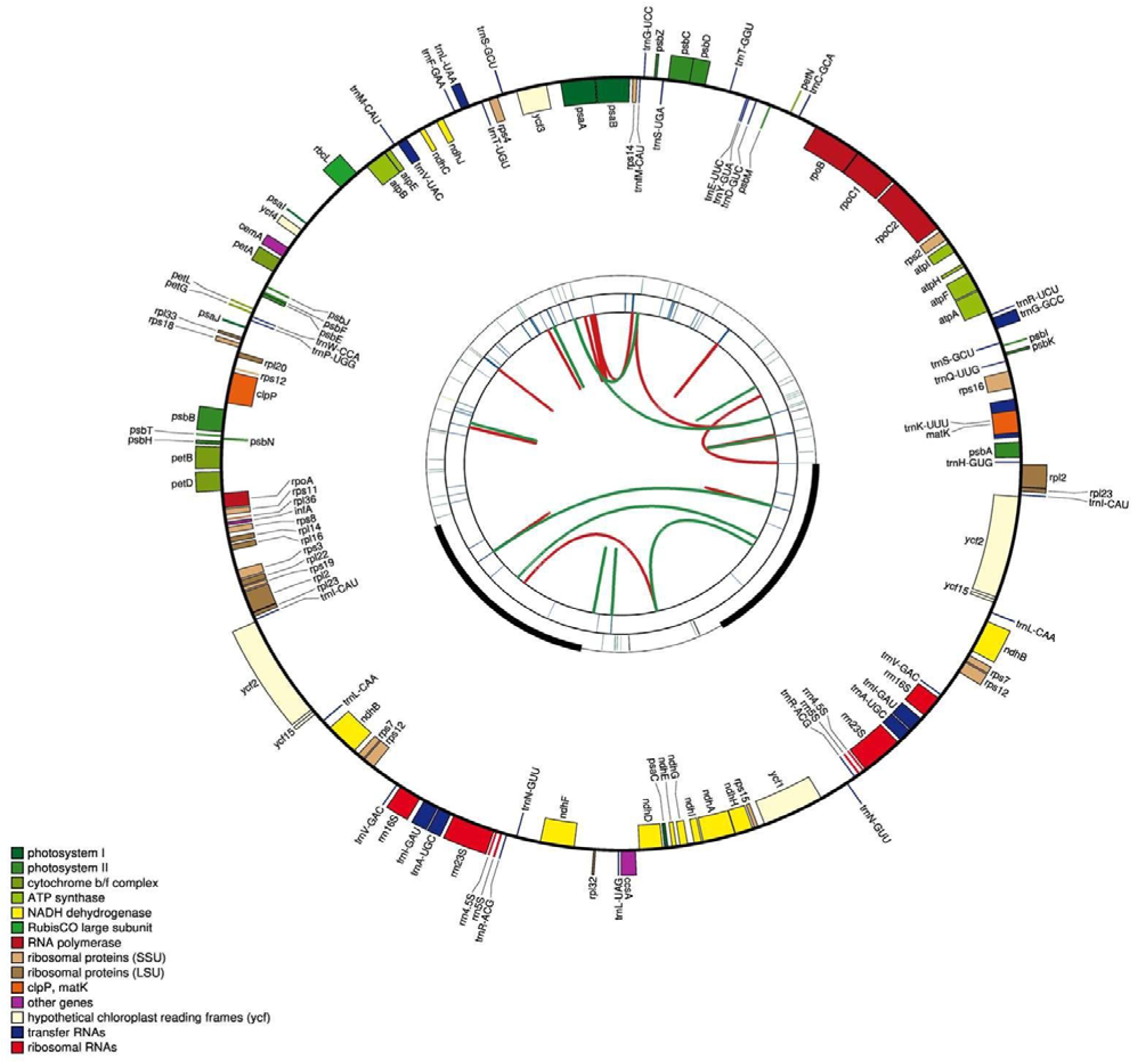
Gene map of the chloroplast genome of D. philippinensis (GB68492). LSC, SSC, and IR regions are indicated. Genes outside the circle are transcribed clockwise; those inside are transcribed counterclockwise.

**Table 1.**
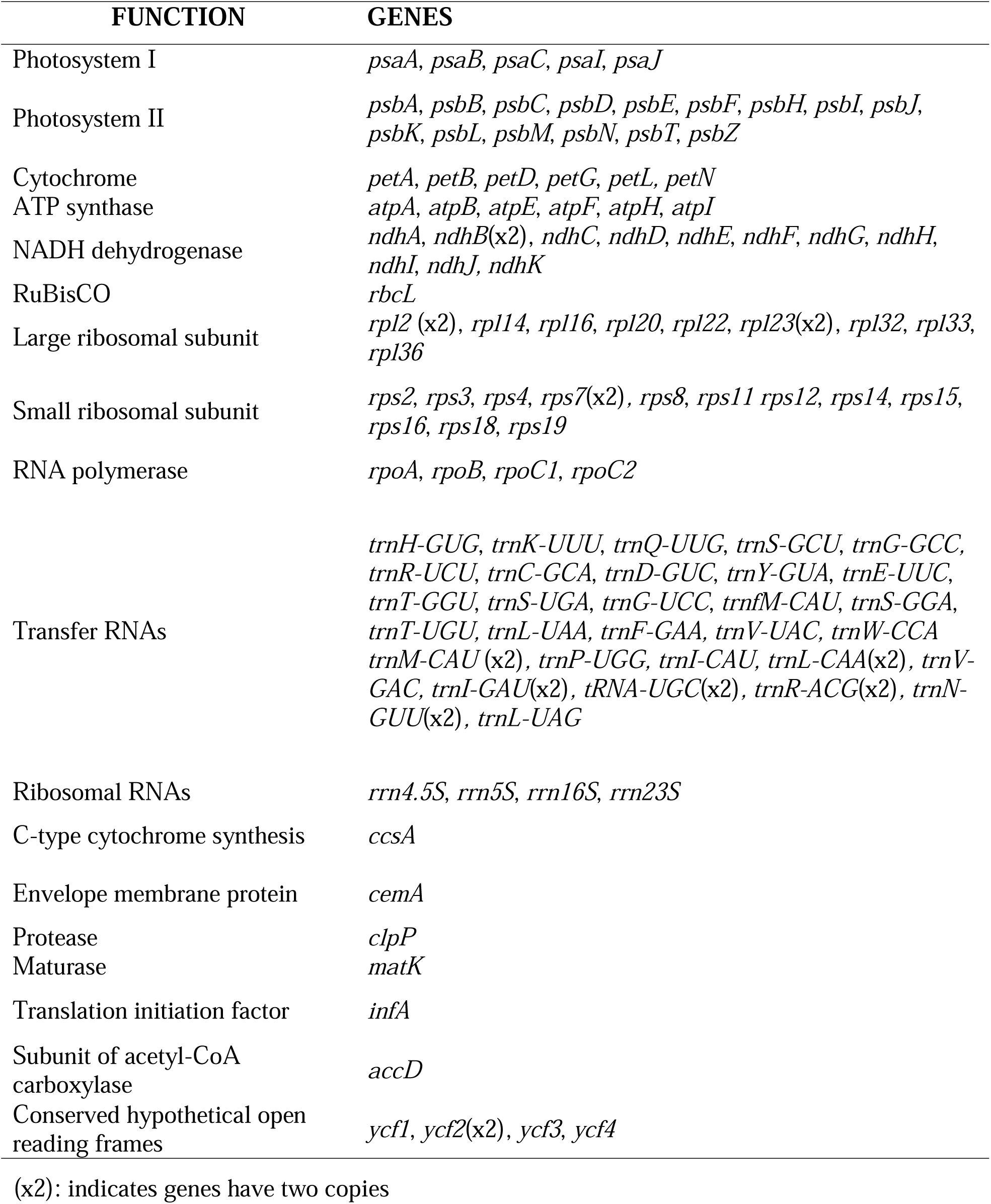
List of genes in the chloroplast genome of D. philippinensis.

Functionally, the genome encodes 79 protein-coding genes, 30 tRNA genes, and 4 rRNA genes, of which 44 are involved in photosynthesis, 59 in self-replication, and 10 in other functions. Initial annotation with CPGAVAS2 predicted 127 genes, while GeSeq identified 135 genes. Discrepancies arose from differences in annotation algorithms, with GeSeq uniquely detecting psbL, rps19, ndhK, ycf1, and accD. BLAST verification confirmed these as functional genes. Conversely, ycf15 was annotated as protein-coding by CPGAVAS2 but is considered non-coding in previous studies (Schmitz-Linneweber *et al*., 2001; Shi *et al*., 2013). After cross-validation, the finalized annotation included 130 genes. This emphasizes that single-tool annotation may miss genes, highlighting the need for multi-tool validation to ensure accurate plastome characterization.

### SSR and Codon Usage Analysis

The chloroplast genome *of D. philippinensis* contained 54 simple sequence repeats (SSRs), 94.4% of which were A/T mononucleotide motifs, confirming a strong AT bias consistent with previous reports (de Souza *et al*., 2019). Dinucleotide (AT/TA) and trinucleotide (ATA) repeats were rare, each occurring once. A total of 53,863 codons were identified, with leucine the most abundant (9.7%, predominantly UUA) and cysteine the least (2.3%, mainly UGU), reflecting typical codon usage bias in chloroplast genomes (Nakamura & Sugiura, 2007).

### Comparative Genome and IR Junctions

Comparison with *D. indica* and *D. turbinata* revealed minor differences in LSC (88,305–90,907 bp) and SSC (18,047–19,349 bp) lengths, while IR regions were nearly identical (Figure 2). No G/C SSR motifs were detected, reinforcing the AT-rich nature of the plastome (Guo *et al*., 2021). IR boundaries were generally conserved, with a slight IRb expansion in *D. indica* due to the rps19 gene.

**Figure 2.**
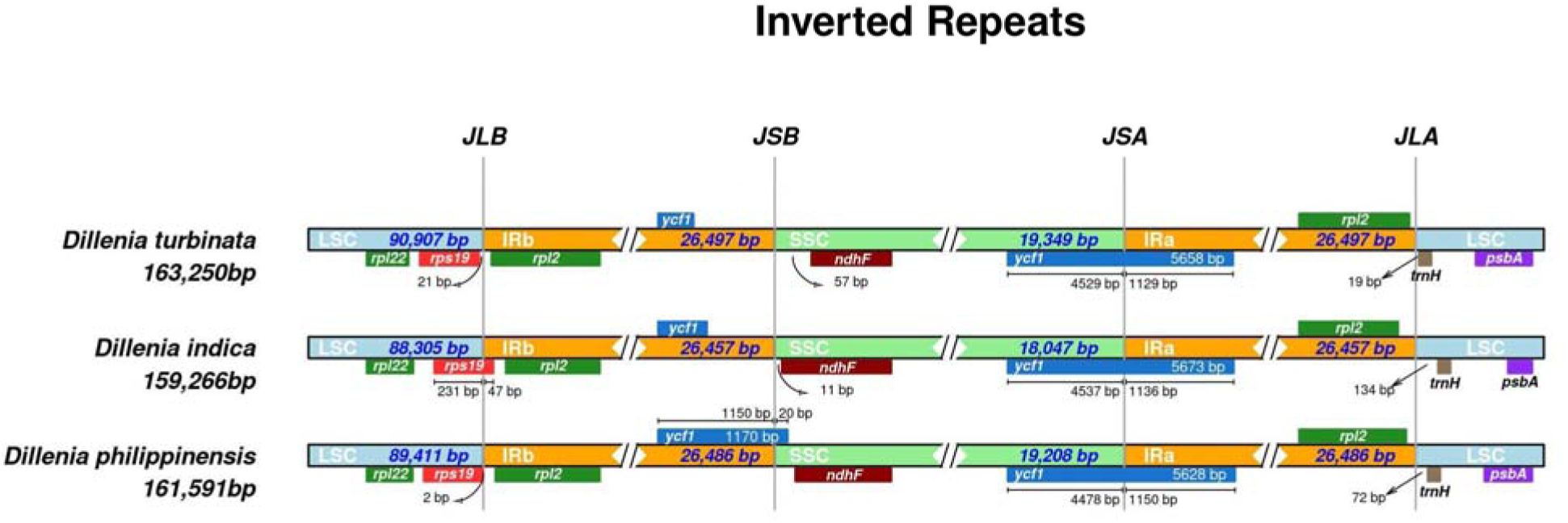
Comparison of IR junctions among D. philippinensis, D. indica, and D. turbinata. JLB: LSC-IRb; JSB: IRb-SSC; JSA: SSC-IRa; JLA: IRa-LSC.

**Figure 3.**
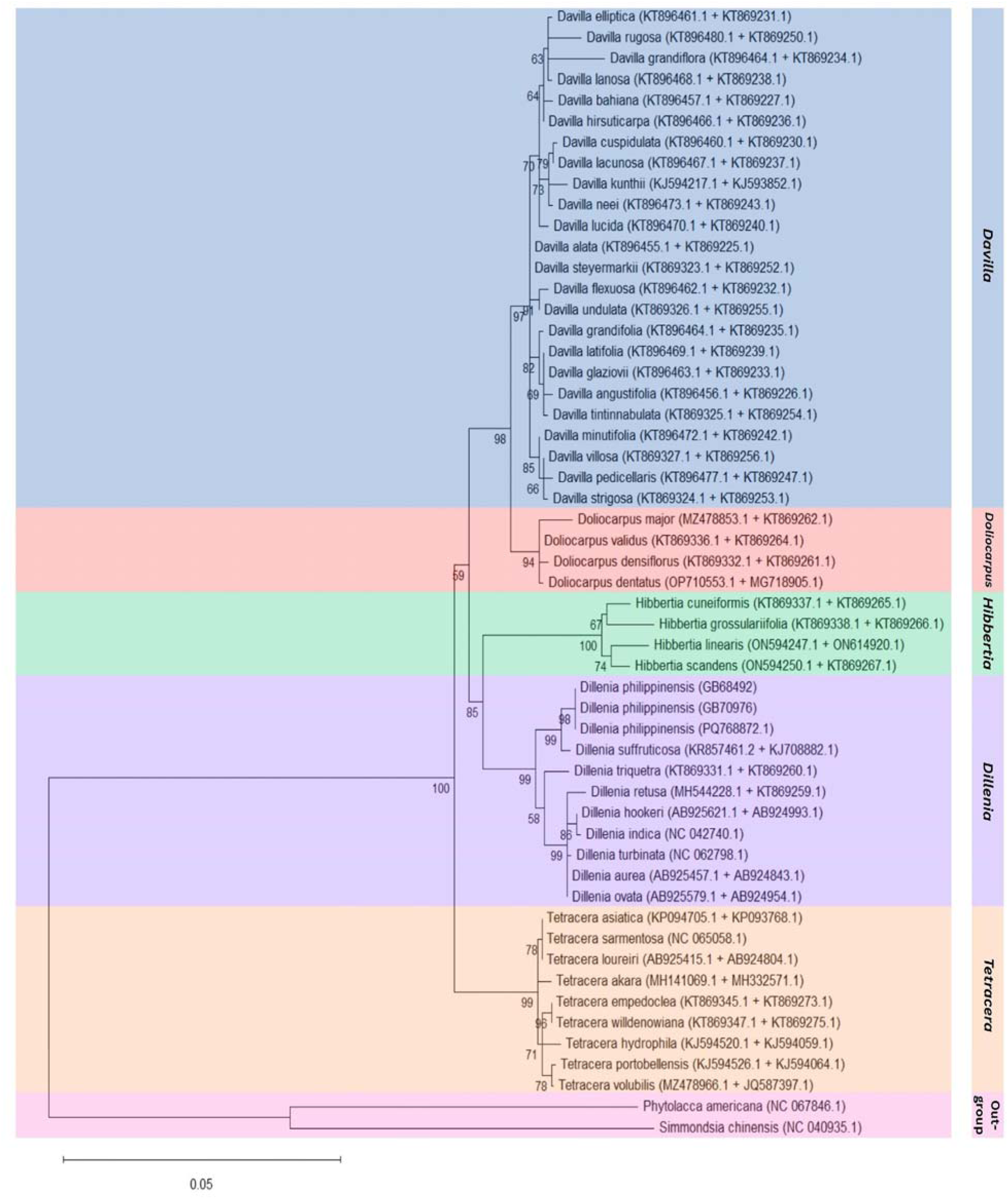
Maximum likelihood phylogenetic tree based on concatenated rbcL + matK sequences, showing D. philippinensis closely related to D. suffroticosa.

### Phylogenetic Analysis

The utility of the assembled chloroplast genome of *D. philippinensis* is limited by the available sequences deposited in NCBI. Currently, there are only three species belonging to the family Dilleniaceae with publicly available complete chloroplast genomes. As such, *the rbcL* and *matK* genes were used to conduct phylogenetic analysis, allowing for the representation of more taxa.

The relationships shown in the tree generated from rbcL, matK, and concatenated rbcL and matK sequences. The concatenated sequences provided higher resolution than single genes (not shown in this paper). It also resolved the discrepancy observed in the trees constructed from individual genes. Tetracera, which was previously depicted to be closely related to *Dillenia* and *Hibbertia*, was depicted to be a sister to the rest of the family, branching off earliest as the closest relative to all remaining members. This result was congruent with those of Horn (2009), who used four plastid loci (*rbcL, infA, rps4,* and the *rpl16* intron). Gontcharov *et al*. (2004) noted that using combined genes in phylogenetic analysis offers greater resolution than single genes and resolves conflicts observed in phylogenetic analyses involving single genes.

The chloroplast genome of *D. philippinensis* provides a foundational resource for species identification, phylogenomics, and conservation of Philippine endemic and indigenous fruits. SSR markers and codon usage patterns can support future population genetics and breeding efforts to enhance the utilization of underexploited species like katmon.

## CONCLUSION

This study reports a complete chloroplast genome of *Dillenia philippinensis* (katmon), a Philippine endemic fruit species. The genome is 161,591 bp with 36.3% GC content and exhibits a typical quadripartite structure comprising 113 unique genes. SSR analysis revealed an A/T-rich repeat pattern, while codon usage analysis indicated a bias toward leucine codons. Comparative analysis with other Dillenia species demonstrated conserved IR boundaries and minor length variations in LSC and SSC regions. Phylogenetic reconstruction based on concatenated sequences of *rbcL* and *matK* consistently placed *D. philippinensis* as closely related to *D. suffroticosa*. These findings provide a foundational genomic resource for germplasm characterization, phylogenomic studies, and the conservation and potential breeding of underutilized Philippine fruit crops.

## Acknowledgment

The authors sincerely thank the National Plant Genetic Resources Laboratory and the Institute of Plant Breeding, University of the Philippines Los Baños, for providing access to plant materials and laboratory facilities essential to this study. We also extend our gratitude to the Philippine Genome Center – Program for Agriculture, Livestock, Fisheries, and Forestry for their technical support and valuable insights throughout the project, and to the Philippine Council for Health Research and Development (PCHRD) of the Department of Science and Technology for funding this research.

We likewise appreciate the colleagues and staff who assisted in sample collection, DNA extraction, and data processing. Their dedication and cooperation greatly contributed to the successful completion of this work.

We express our heartfelt gratitude to our mentors, families, and peers for their encouragement and unwavering support. Finally, we extend our deepest appreciation to the Filipino people whose contributions as taxpayers made this research possible; this work is dedicated to them.

## Disclaimers

The views and conclusions expressed in this article are solely those of the authors and do not necessarily represent the views of their affiliated institutions. The authors are responsible for the accuracy and completeness of the information provided, but do not accept any liability for any direct or indirect losses resulting from the use of this content.

